# Modeling Spatial Genomic Interactions with the Hawkes model

**DOI:** 10.1101/214874

**Authors:** Anna Bonnet, Vincent Rivoirard, Franck Picard

## Abstract

The spatial localization of many DNA-protein interactions is now available thanks to the development of ChIP-Seq, and their investigation calls for adapted statistical methods. Many methods were developped for peak calling, but few were proposed for the downstream analysis of peak-like data, whereas the spatial structure of such data may contain relevant biological information, like binding constraints for instance. Associations between the occurrences of two genomic features are usually assessed by overlaps, but here we propose a statistical model to precisely quantify the spatial interactions between the location of binding events. Our methodology relies on a multivariate spatial process, the Hawkes model, that can also be interpreted in terms of a graphical model to highlightspatial dependencies between genomic features. Using our method, we explore the chromatin landscape of replication origins, and we highlight attractive and repulsive patterns that can be related to the regulation of the spatial program of replication. We compete our method with both pairwise and multivariate approaches, implemented in the packages **`GenometriCorr`** and **`ppstat`** respectively. We show that our procedure describes with more details the complex patterns of spatial interactions and also provides estimates that are very convenient for interpretation.

## 1 Introduction

Next generation sequencing technologies allow to study genomes with an unprecedented resolution; molecular processes can be captured through the mapping of the whole genetic variants of a genome, the measure of the expression of all genes of a cell, and now the structure of chromatin and its modifications. Epigenetics has certainly been one of the most active fields for the last few years, thanks to the development of ChIP-Seq that allow the spatial localization of DNA-proteins interactions. One of the major challenges now is to extract relevant information from this amount of data, to better understand genomes regulation.

ChIP-Seq data provide maps of chromatin modifications along chromosomes, and we focus on the statistical modeling of this spatial specificity of the data. A first category of methods is based on Hidden Markov Models (HMMs) and aims at detecting regions in the ChIP-Seq signal. The ChromHMM method (Ernst and Kellis, 2012) detects from the raw signal the presence or absence of each chromatin mark and segments the genome in regions characterized by specific combinations of marks. This segmentation defines a “chromatin landscape”, which describes the modifications of chromatin states and their associated biological functions. Wu and Zhaohui (2013) developed the GiClust multivariate procedure, also based on HMMs, that uses ChIP-seq data after peak detection. GiClust provides a segmentation in clusters on which the probability of occurrence for each feature is estimated.

Several methods were also developed to study the co-occurrences of a group of features by testing their pairwise associations. These methods are very general and provide co-occurrences patterns for any spatial data described by points or intervals, in particular ChIP-Seq data after any peak detection method. Favorov et al. (2012) propose four standard methodologies (relative distance test, absolute distance test, projection test and Jaccard test), all implemented in the GenometriCorr package. D. Chikina and G. Troyanskaya (2012) noticed that most procedures used to compare sets of intervals considered the binary situation-with or without overlaps- and defined a metrics to assess the distance between two intervals lists. We will show that this package provides results that may not be adapted to the multivariate framework, because the pairwise strategy does not account for spurious correlations, and calls for multiple testing procedures that are not provided by the package. Finally, Wei and Wu (2016) proposed to use the ChromHMM segmentation to account for genome heterogeneities before looking for associations between genomic features.

In this work we focus on the modeling of spatial interactions between called ChIP-Seq peaks, and more generally between maps of genomic intervals structured along chromosomes. We propose a modeling based on point patterns to account for potential physical interactions that come from the spatial proximity of binding events. To propose a fine-scale modeling of spatial interactions, we focus on the spatial covariance between points, embedded within a multivariate model. This multivariate aspect of our model is central compared to existing pairwise strategies, since it allows to correct for spurious aliasing correlations. Then the results of our method can be interpreted in terms of spatial attraction/repulsion between genomic features, which could help in the identification of characteristic distances for instance. Our strategy is model based, and relies on the modeling of spatial interactions with the Hawkes model, for which non parametric estimation procedures were recently proposed by Reynaud-Bouret et al. (2014). The benefits of this method compared to standard approaches are plural, in particular it allows the reconstruction of sparse interaction functions, which is very convenient for interpretation. In particular, this sparse method produces much more interpretable interaction functions, as compared with spline-based estimates. Moreover, based on these functions, one can reconstruct a graphical model with an edge for each significant interaction between pairs of features. Let us also note that defining a model could also be useful to generate realistic spatially interacting genomic features.

This method is used to explore the spatial interaction landscape of replication origins with their chromatin context. DNA replication is a biological process that is intrinsically spatial and the identification of replication origins at fine scales has revealed complex interplays with genetic and epigenetic features in the early steps of DNA replication (Picard et al., 2014; Julienne et al., 2015). Using our method, we reveal attractive effects between marks and origins, with estimated intensities and distances of interaction. We also show strong attractions and repulsions between marks, which confirms the necessity of including all marks in a multivariate model.

## 2 Model and method

### 2.1 Multivariate Hawkes process

We consider a dataset with *M* maps of ChIP-Seq peaks, or more generally of mapped genomic features. We denote by *X_m_* the *m*-th map, such that 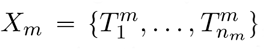 with 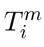 the genomic coordinate of the peak *i* in map *m*. In a first approximation, we reduce the ChlP-Seq intervals to single points. Then, we consider that these positions are random to account for biological and technical noise and we model (*X*_1_,…, *X_M_*) as a multivariate point process. To model the spatial interactions between genomic features, we introduce the conditional intensity of the process, which models the probability of occurrence of points, such that: for 1 ≤*m* ≤ *M*, for *t* ∈ [*T*_1_, *T*_2_] a given genomic region,

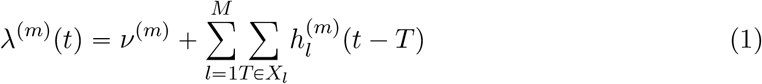

- 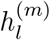 is a function defined on (0, +∞) that characterizes the influence of each occurrence *T* ∈ *X_l_* on *T*′ ∈ *X_m_*. For instance, if 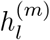 is a positive function on (0, *d*) and is null elsewhere, the probability of observing points of *X_m_* will increase within a distance of *d* after observing points of *X_i_*.
- *v*^(*m*)^ is a spontaneous rate, it corresponds to the part of the intensity *λ*^(*m*)^(*t*) that cannot be explained by the occurrences of (*X*_1_,…,*X*_M_).

*Remark* 1. The function 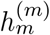 describes the self-interactions between the occurrences of the *m*th process. In particular, it can model regions with clusters of occurrences of the same process.

*Remark* 2. In Model (1), the interaction functions 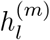 and 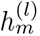 can be (and generally are) different, so that we can differentiate the effect of *X_m_* on *X_l_* from the one of *X_l_* on *X_m_*. Model (1) can be interpreted as a directed graphical model with M nodes and edges for each non-zero interaction function.

Our goal is to propose an estimator of(*v^(m)^*, 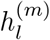)_*l,m*_) in order to understand the spatial interactions between all the point processes.

### 2.2 Method

Reynaud-Bouret et al. (2014) proposed a methodology to estimate the interaction functions, a brief description of which is given in this section. A more detailed version is given in the Appendix, section 5. The procedure is nonparametric, which means that there is no assumption on the form or regularity of the intensity functions that we want to estimate.

The main idea is to find a candidate 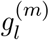 to estimate 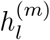 that can be decomposed on a histogram basis as follows:

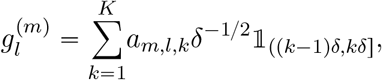

where *δ* is the size of each bin and *K* the number of bins. The product *Kδ* corresponds to the maximal distance between two occurrences that interact with each other. The coefficients *a*_*m*,*l*,*k*_ can be interpreted is terms of spatial covariance between points of *X_l_* and points of *X_m_* at lag included in ((*k* — 1)δ, *kδ*], corrected by other potential aliasing covariates.

As detailed in the Appendix, we propose to estimate the coefficients (*a*_*m*,*l*,*k*_)_*l*,*m*,*k*_ using a penalized least squares criterion with theoretically calibrated weights, which provides a sparse estimation of the interaction functions 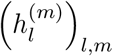. This constitutes an alternative to other proposed strategies based on splines Carstensen et al. (2010) that create wiggles that are difficult to interpret in practice, and that is less powerful to detect small scale interactions, as shown in the next section.

## 3 Application

### 3.1 Data

The spatio-temporal program of replication is tighly regulated in cells, as it allows the faithful duplication of the genetic material. Thanks to the recent identification of replication origins (Picard et al., 2014), much progress has been made in the identification of the genetic and epigenetic basis of these controls (Picard et al., 2014; Julienne et al., 2015). Current studies proposed to unravel the interactions between replication origins and chromatin marks using standard associations, by studying the overlap between genomic maps. However, the precise spatial interactions between origins and surrounding marks have never been investigated for lack of appropriate method. The data we study here are the positions of replication origins as well as the positions of five chromatin marks: H3K9ac, H3K4me3, H3K27me3, H3K9me3 and H4K20me1, for the HeLa cell line. We deal separately with regions replicated early and late in the cell cycle, as regulation mechanisms were suggested to be different (Picard et al., 2014).

### 3.2 Results

In the following we display the estimated interaction functions, with positive and negative support whereas Model (1) only considers positive support (backward interactions). Forward interactions were quantified by running the method on the same data but in reverse order. We compete our method with the method of Carstensen et al. (2010) that is also based on the Hawkes model, but that uses B-splines to estimate the interaction functions. To proceed we used the **`ppstat`** package with the settings proposed in Carstensen et al. (2010). Note that the number of splines knots (8) is comparable to K in our model.

#### 3.2.1 Influence of the marks on origins

We start to investigate the influence of chromatin marks on origins with the pairwise associations as proposed by the `GenometriCorr` package (Table 1). Four statistics are proposed (Jaccard index, absolute distance, relative distance and projection) to assess significance of the proximity between each mark and origins. Unfortunately, the raw p-values are not accessible, and no multiple testing procedure is included in the package, whereas the multivariate framework calls for a control of false positive associations. Moreover, the pairwise framework does not account for spurious correlations (contrary to our procedure for instance). Consequently, nearly all tests of associations are significant (average distance smaller than expected and average overlap larger than expected under independence), except the projection test for H3K9me3 in early regions (which is consistent with the null interaction that we estimate). Thus `GenometriCorr` provides far less informative results regarding the multivariate framework and the precise shape of the spatial interactions between marks and origins.

Thanks to our probabilistic framework, we estimate interaction functions as displayed in Figure 1 to quantify the influence of each mark on the presence of origins. One main advantage of the histogram-based lasso estimate we propose is that the estimated value of 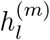 on the interval [(*k* — 1)*δ*, *kδ*] is the mean number of additional points of *X_m_* due to the presence of each point of *X_l_* Hence, our results show that each occurrence of H3K9ac within a distance of 1 kb of origins induces an average increase of ∼ 4.5 origins in early regions, and ∼ 6 origins in late regions. Our results show that the presence of most of the marks increases the number of origins in both early and late regions, within a small distance (< 3kb, except for H3K27me3, whose effect is more diffuse). In particular, the influence of H3K9me3 is specific to late regions, which is consistent with the correlation study of Picard et al. (2014). However, the effect of the timing is less noticeable for the other marks, which might be explained by our division of regions between two groups (early and late) while Picard et al. (2014) compared the associations between origins and marks divided in 6 timing groups.

**Figure 1.**
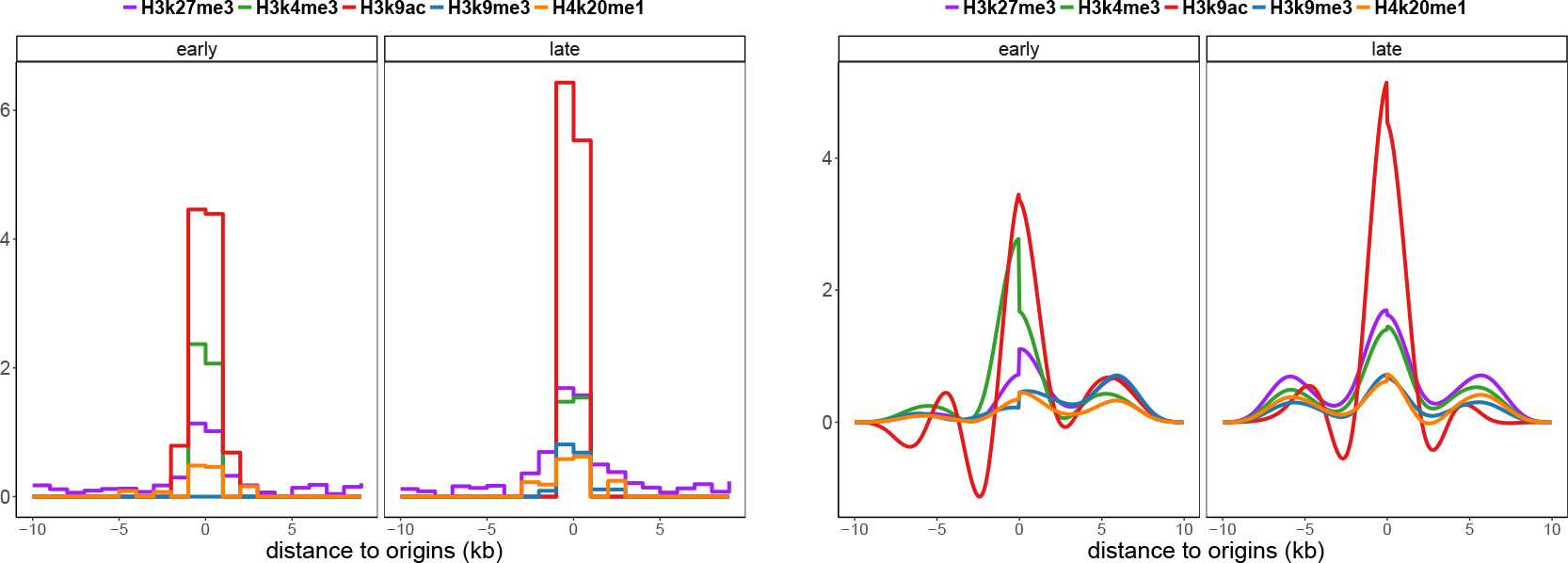
Estimated interaction functions 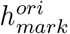 for all marks according to timing categories (early/late). Left: lasso estimate with δ = 1kb and *K* = 10, right: splines estimates.

When compared with the results obtained with the **`ppstat`** package, the interactions functions show similar trends, but splines induce wiggles that are both positive and negative, which makes these interaction functions less interpretable. Moreover, from the computational point of view, **`ppstat`** can not handle the full genomic datasets, and we had to split the data into subsets with averaged results. Note that the spline framework provides confidence intervals that are not displayed in the Figures.

**Table 1.**
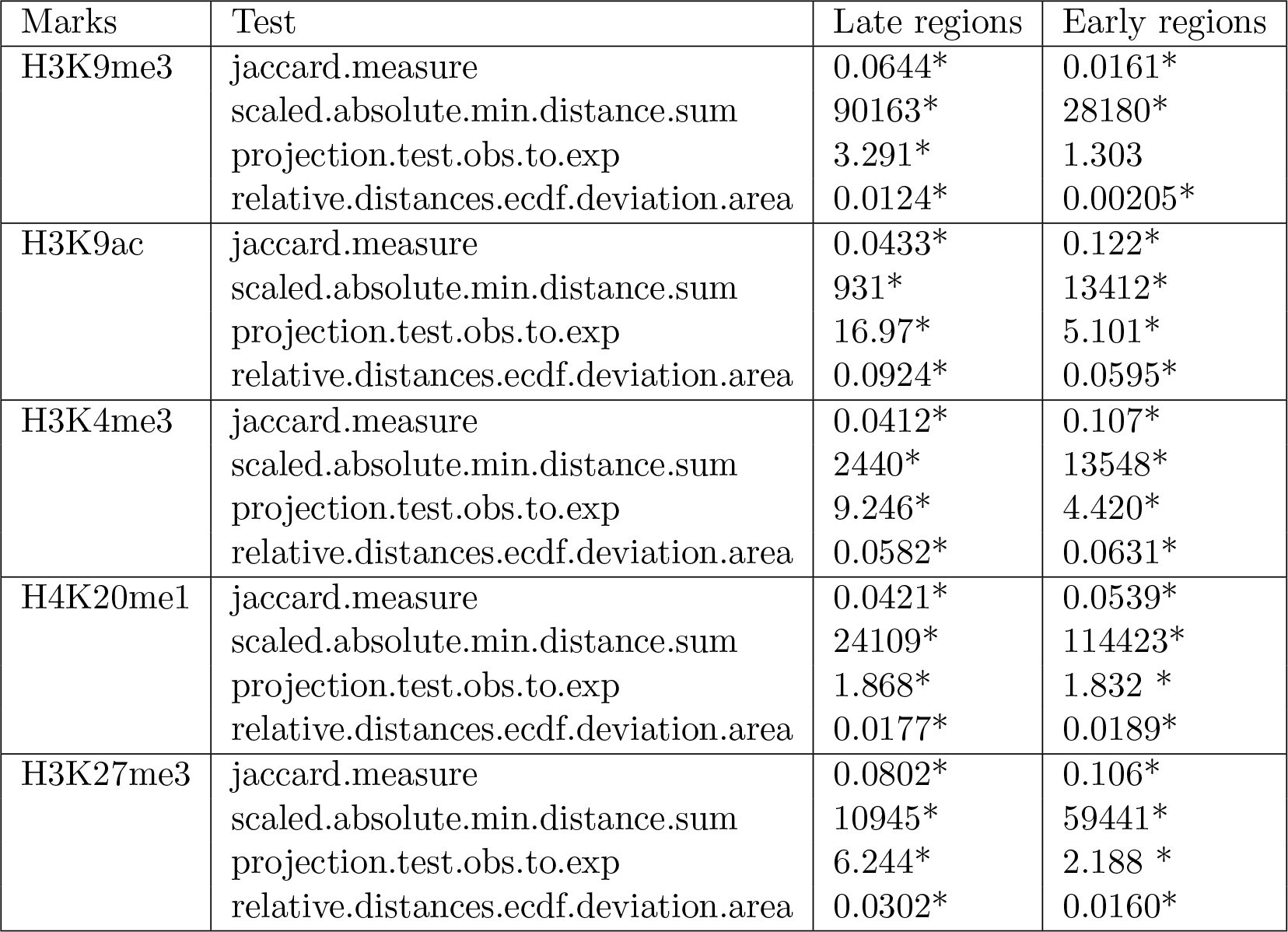
Associations between marks and origins detected by the `GenometriCorr` package. The star next to the test statistics indicates that the corresponding test is significant, with a p-value smaller than 0.01.

#### 3.2.2 Interaction between marks

Figure 2 shows the interactions between chromatin marks. We observe a very strong attraction between H3K4me3 and H3K9ac (2nd column) which are both open-chromatin marks. These marks have also attractive interactions, though weaker, with H4K20me1. Our results show that these three marks have a repulsive interaction with H3K27me3 (1rst column), although they all have an attractive effect on origins. Finally, H3K9me3 is also repulsed by the presence of the other chromatin marks since its interaction functions are all negative (the intensity of repulsion is stronger in early). Finally the landscape of H4K20me1 is dynamic between early and late: while H3K27me3 is a strongly repulsive in a ∼10kb range, whereas H3K9me3 becomes strongly repulsive in late regions. These different behaviors are not detected when using splines (Fig. 5).

**Figure 2.**
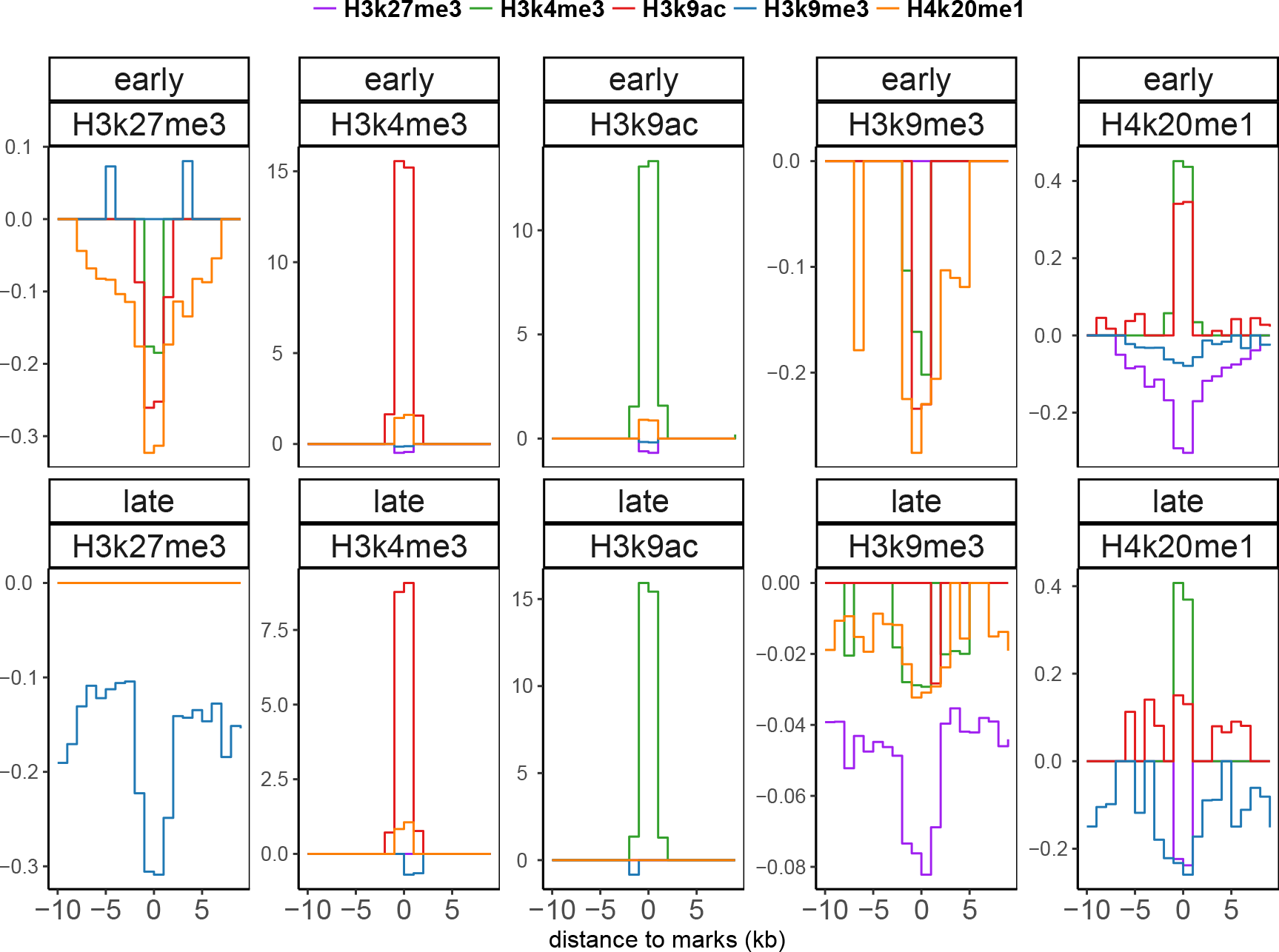
Lasso estimates of interaction functions between marks, with δ = 1kb and *K* = 10.

**Figure 3.**
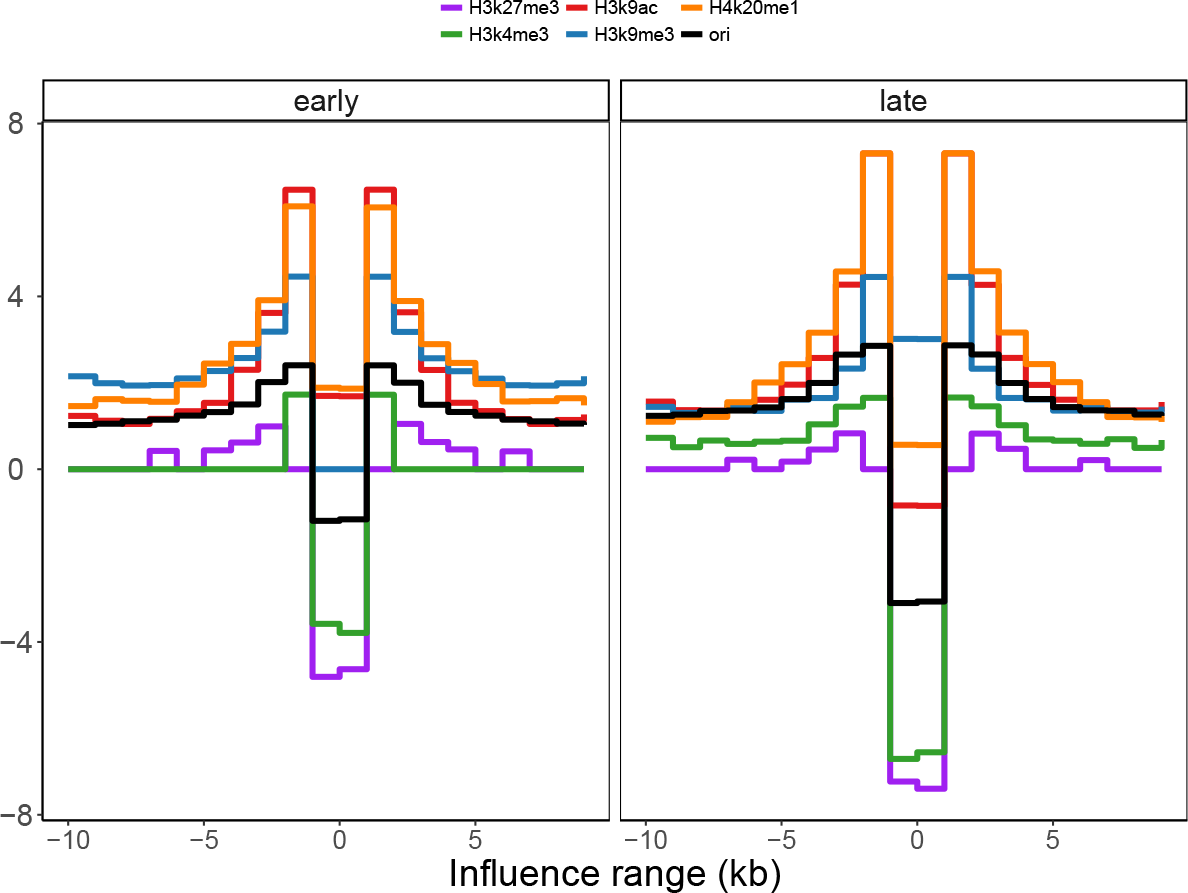
Lasso estimates of self interaction functions, with δ = 1kb and *K* = 10.

**Figure 4.**
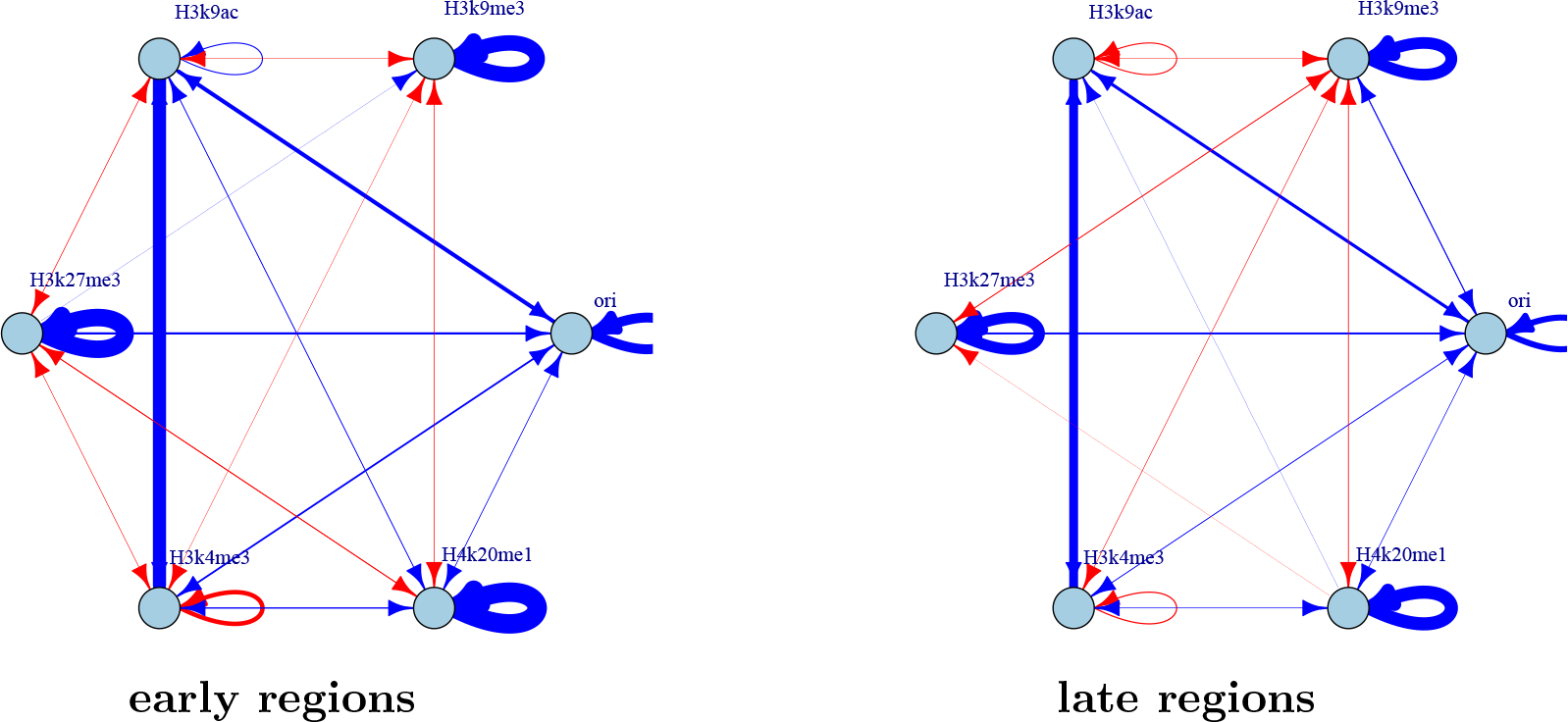
Graph of interactions between marks or origins. The repulsive effects are in red, the attractive effects in blue. The width of arrows is proportional to the integrated absolute value of interaction function.

**Figure 5.**
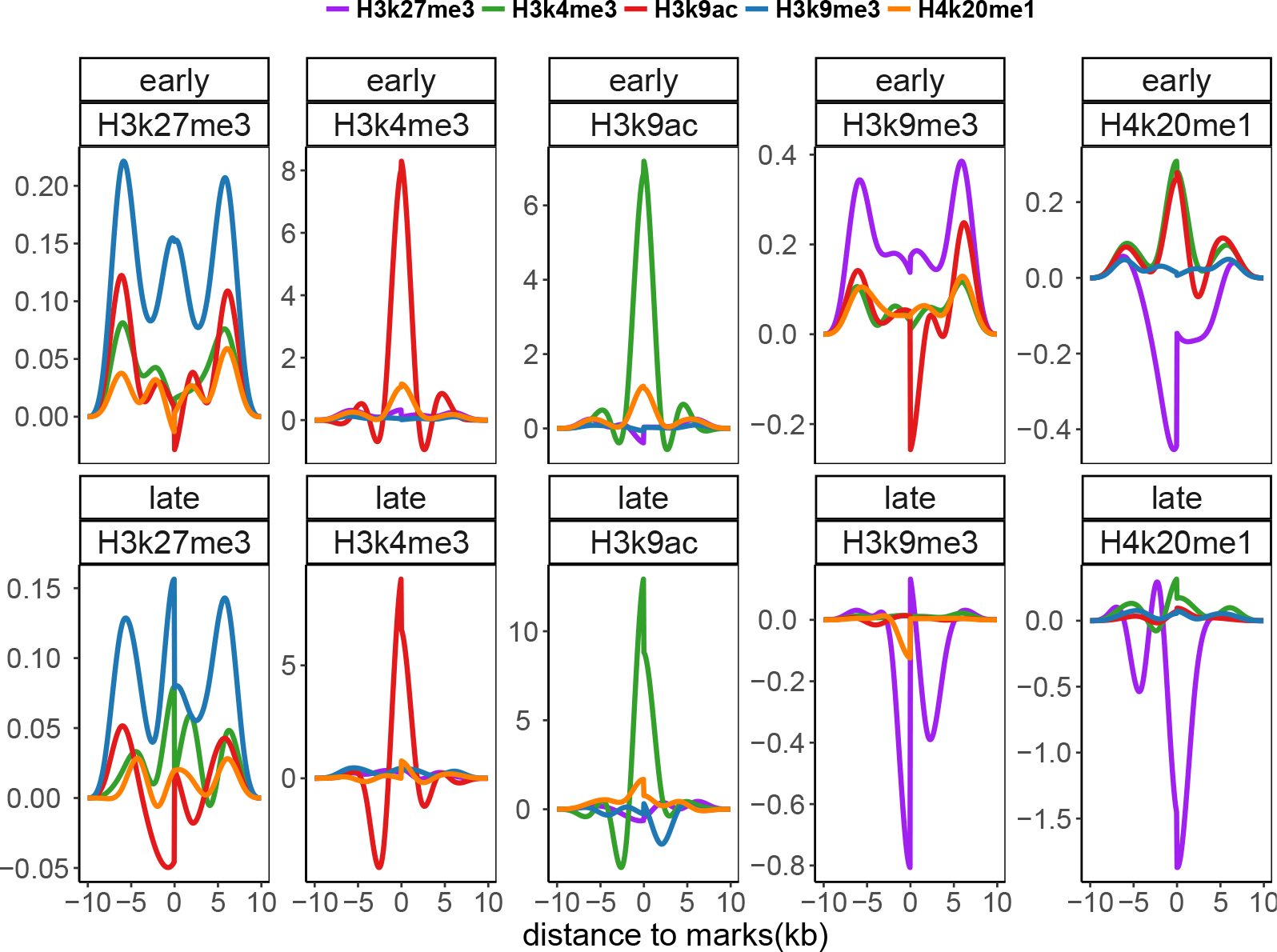
Splines estimates of interaction functions between marks.

As mentioned in Section 1, the multivariate approach that we use allows us to correct the pairwise interactions by taking into account the interactions with other features. Here, we see several interactions between marks which are likely to impact the associations mark/origins that we might detect without this correction.

#### 3.2.3 Self interactions

Figure 3 shows the estimated 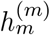 functions of Model (1), that is the self interactions between occurrences of origins, and between the occurrences of each mark. Our results show that self interactions are strong for origins and for most marks, which suggests that there are regions with clusters of points. Note that that self interactions are weaker for the two open-chromatin marks H3K4me3 and H3K9ac. Let us also observe that for origins and several marks, the estimated 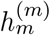 take negative values on [-1,1kb] which is due to the modeling where we consider our observations as points when they are actually centers of intervals; the distance between two centers is indeed obviously at least larger than the length of the intervals. Note that this feature is smoothed away when using splines (Fig. 6).

**Figure 6.**
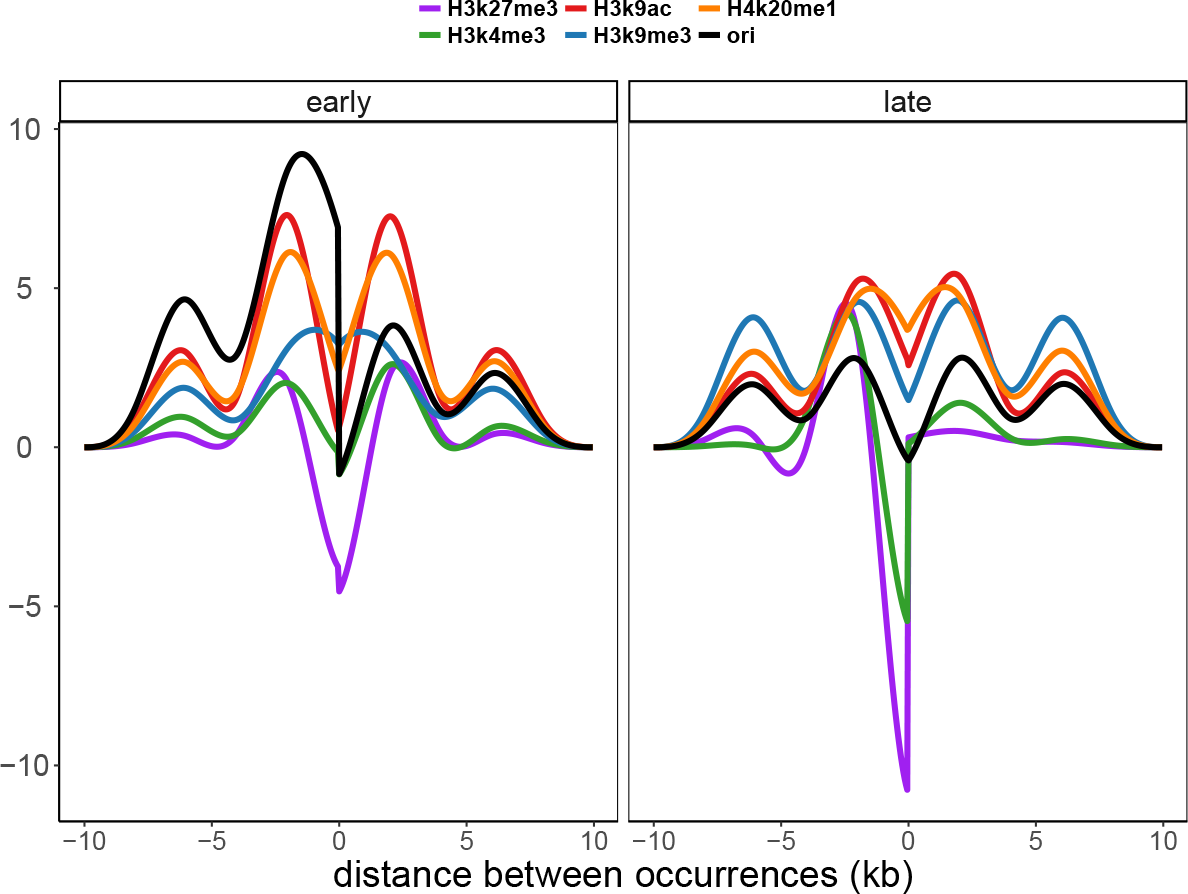
Splines estimates of self interaction functions

#### 3.2.4 Interaction graph

The estimated interactions are summarized in a graph (Figure 4), the edges of which have a width proportional to the integrated interaction function between two nodes. It confirms that there is groups of marks composed of H3K4me3, H3K9ac and H4K20me1 that have attractive effects on each other and on origins, and repulsive effects on the other marks. H3K27me3 and H3K9me3 attract only origins but repulse the other marks (with only a small attraction detected between H3K27me3 and H3K9me3 in early regions). Although the differences between the early and late graphs are quite small, we notice that the edge between H3K9me3 and origins appears only in the late graph.

## 4 Discussion

We propose a probabilistic model to quantify spatial interactions between genomic features, with application to the epigenetic landscape of replication origins. Our method allows us quantify these interactions with much more precision than classical correlation or overlaps approaches. Moreover, the sparse Lasso estimate provides more interpretable results compared with splines. Our model has revealed several attraction/repulsion patterns with typical distances of interaction, and we show that these patterns are dynamic between early and late regions. Note that these patterns are based on statistical correlations, which may only reflect one aspect of the complex epigenetic landscape of replication origins. The statistical comparison on interaction functions between timing regions is an important perspective of our work. Strong statistical developments will be needed to achieve this goal, which can be rephrased as a two-sample test for Hawkes processes, and which will be based on non parametric testing to compare intensity functions. Finally, the interpretation of the Hawkes model as a graphical model allows us to represent our result in a very synthetic graph. Let us also note that the methodology we propose can be generalized to any set of spatially ordered genomic features. The next step forward will also be to enrich our method by accounting for the variability in the detection of the genomic features.

## 5 Appendix

We give in this section more details on the estimation method proposed by Reynaud-Bouret et al. (2014).

Let us denote 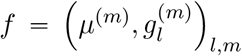 a candidate to estimate 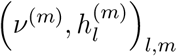 and the corresponding estimator of λ^(*m*)^:

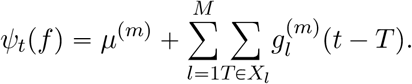

We want the intensity candidates to minimize the *ℓ*_2_-distance to the intensites *λ*^(1)^,…, λ^(*M*)^:

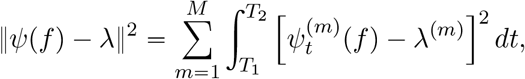

which is equivalent to minimizing the contrast 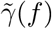 defined by

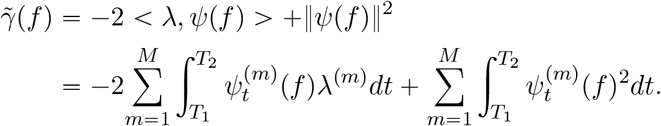

Since a natural approximation of λ^*(m)*^(*t*) is the point measure *dN*^*(m)*^ (*t*), we shall focus on minimizing γ(*f*) defined by

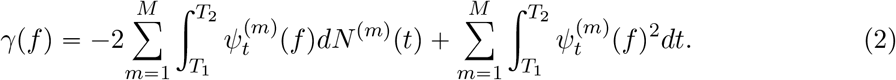

The main idea is to find a candidate 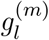 to estimate 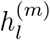 that can be decomposed on a histogram basis as follows:

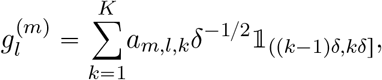

where *δ* is the size of each bin and *K* the number of bins. The product *Kδ* corresponds to the maximal distance between two occurrences that interact with each other. The coefficients *a*_*m*,*l*,*k*_ can be interpreted is terms of spatial covariance between points of *X_l_* and points of *X_m_* at lag included in ((*k* — 1)δ, *kδ*]. Then, if we use this decomposition in the definition of ψ_*t*_(*f*),

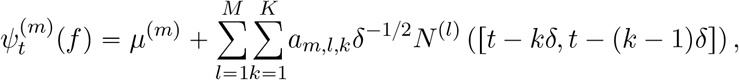

where *N*^(*l*)^(^*I*^) denotes the number of points of *X_l_* on interval *I*. If we replace 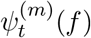 in (2), Reynaud-Bouret et al. (2014) show that the contrast can be rewritten as:

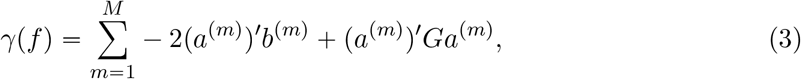

where
(*a*^(*m*)^) = (μ^(*m*)^, *a*_*m*,1,1_,…, *a*_*m*,1,*K*_,*a*_*m*,2,1_,…, *a_m_*,*M*,*K*) are the coefficients to estimate, and *b^(m)^* and *G* are defined by:

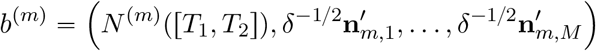

where

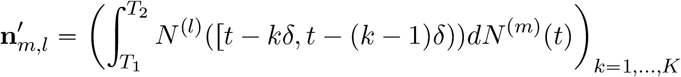

and

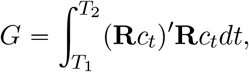

where

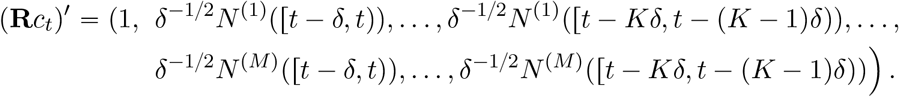

The solution of the minimization of (3) can be easily obtained; in particular, if *G* is invertible

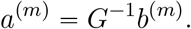

Notice that the entries of *b^(m)^* correspond to pairwise correlograms, while the coefficients of *a^(m)^* are corrected by other potential aliasing covariates, the effects of which are quantified by the matrix *G*.

The procedure is achieved by thresholding the coefficients of *a^(m)^*, as follows:

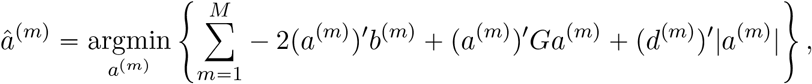

where the weights (*d^(m)^*)1≥*m*≥*M* are theoretically calibrated. Once the support is determined by the previous procedure, the final estimation of the non-zero coefficients of *a^(m)^* is obtained thanks to a classical least-squares estimate on this support. This two-step approach allows to overcome the bias issue of the LASSO estimates.

## References

Carstensen, L., A. Sandelin, O. Winther, and N. R. Hansen (2010, Sep). Multivariate hawkes process models of the occurrence of regulatory elements. BMC Bioinformatics 11(1), 456.

D. Chikina, M. and O. G. Troyanskaya (2012). An effective statistical evaluation of chipseq dataset similarity. Bioinformatics 28(5), 607–613.

Ernst, J. and M. Kellis (2012, March). Chromhmm: automating chromatin-state discovery and characterization. Nat Meth 9(3), 215–216.

Favorov, A. V., L. Mularoni, L. M. Cope, Y. A. Medvedeva, A. A. Mironov, V. J. Makeev, and S. J. Wheelan (2012). Exploring massive, genome scale datasets with the genometricorr package. PLoS Computational Biology 8(5).

Julienne, H., Audit B., and A. Arneodo (2015). Embryonic stem cell specific “master” replication origins at the heart of the loss of pluripotency. PLoS Comput Biol 11(2), e1003969.

Picard, F., J.-C. Cadoret, B. Audit, A. Arneodo, A. Alberti, C. Battail, L. Duret, and M.-N. Prioleau (2014). The spatiotemporal program of dna replication is associated with specific combinations of chromatin marks in human cells. PLoS Genet 10(5), e1004282.

Reynaud-Bouret, P., V. Rivoirard, F. Grammont, and C. Tuleau-Malot (2014, April). Goodness-of-fit tests and nonparametric adaptive estimation for spike train analysis. Journal of Mathematical Neuroscience, 4:3.

Wei, Y. and H. Wu (2016). Measuring the spatial correlations of protein binding sites. Bioinformatics 32(12), 1766–1772.

Wu, H. and S. Q. Zhaohui (2013). Exploring the cooccurrence patterns of multiple sets of genomic intervals. BioMed Research International.

